# Acid enhances salt taste by activating the epithelial sodium channel

**DOI:** 10.1101/2023.04.14.536795

**Authors:** Daniel M. Collier, Zerubbabel J. Peterson, Alan J. Ryan, Noah N. Williford, Christopher J. Benson, Peter M. Snyder

## Abstract

Salt is often used to enhance the flavor of foods and drinks, and in turn, some foods and drinks may intensify the taste of salt. For example, highly acidic carbonated beverages sometimes accompany salty snacks, and margaritas made with acidic citrus are served in salt-rimmed glasses. However, whether and how acid might enhance salt taste remain unknown. Epithelial sodium channels (ENaC) in tongue taste cells detect dietary sodium. We found that acid irreversibly increased ENaC channel activity, with half-maximal activation occurring at pH 2.6. Acid altered ENaC gating by increasing the rate of channel opening and reducing the rate of channel closing. Acidic beverages Coca-Cola® and Pepsi® (pH 2.2-2.4) also stimulated ENaC current but Diet Coke® (pH 3.2) did not. In humans, we found that acid reduced the sodium taste detection threshold. These findings identify a functional interplay between dietary sodium and acid--by modulating ENaC gating, acid enhances salt taste.

## Introduction

Dietary sodium is detected in part by the epithelial sodium channel ENaC. A heterotrimer of α, β, and γ subunits, ENaC is expressed in taste cells located in fungiform and foliate papillae of the tongue (*1-3*). Salt taste in the appetitive range (<150 mM) is disrupted by the ENaC blocker amiloride (*4*) and by targeted deletion of αENaC in mouse taste receptor cells (*5*). In addition to its role as a salt taste receptor, ENaC functions as a pathway for urinary sodium reabsorption in the kidney collecting duct and connecting tubule (*6-8*). Mutations that alter ENaC activity disrupt sodium balance, causing inherited forms of hypertension (Liddle’s syndrome) (*9-11*) and hypotension (pseudohypoaldosteronism type 1) (*12*). Thus, by modulating sodium intake and sodium excretion, ENaC is positioned at two critical locations to control sodium balance.

We seldom encounter dietary sodium in isolation—we eat and drink complex mixtures that interact in ways that alter our experience of taste. For example, previous work suggested that acid may modulate salt taste, but the data were conflicting (*13-18*). Moreover, molecular mechanisms to explain these interactions have not been determined; it is not known if they occur at the level of the receptor or at a more central point in the sodium taste transduction pathway.

We previously reported that ENaC is stimulated in a reversible manner by acid within the range found in the kidney collecting duct (pH 8.5-4.5) (*19-21*). However, in the tongue, ENaC is exposed to more extreme changes in pH through the ingestion of acidic foods and beverages. For instance, lime juice has a pH of 1.8 and some carbonated beverages are also highly acidic. Here we asked whether protons in this highly acidic range alter ENaC activity and salt taste.

## Results and Discussion

We quantified ENaC sodium current before and after exposure of the extracellular side of the channel to acid (pH 2.5). Acid produced a transient increase in current (Fig. 1A). Moreover, when the acid was washed away (pH returned to 7.4), current increased to a level greater than the pre-application baseline. Amiloride (an ENaC blocker) abolished this current, indicating that ENaC was responsible. Acid increased ENaC current 2.3-fold (Fig. 1B).

**Fig. 1.**
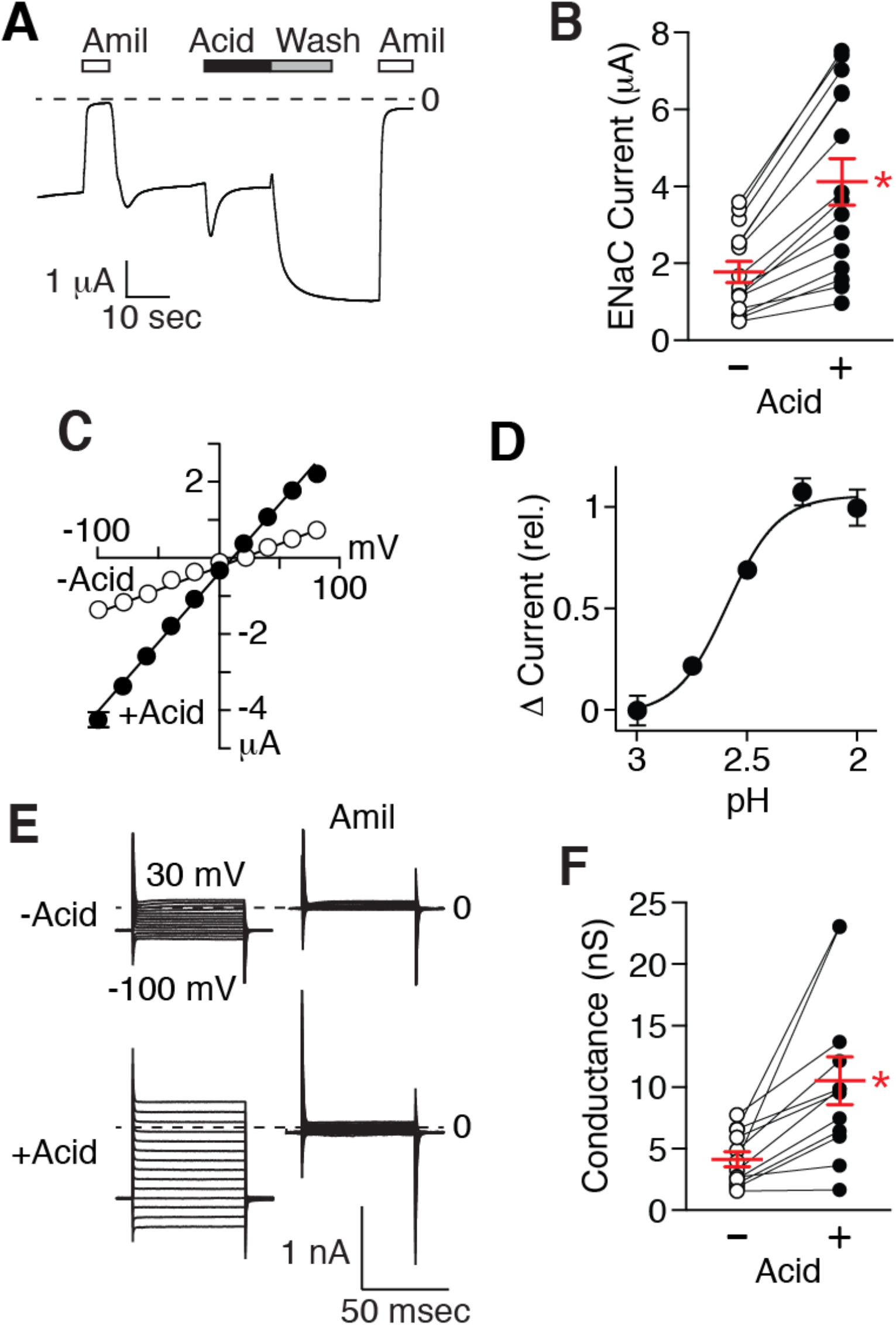
Acid stimulates ENaC current. (**A**) Current trace from a *Xenopus* oocyte expressing human αβ γENaC obtained by two-electrode voltage clamp (holding potential -60 mV). Amiloride (10 μM) and acid (pH 2.5) were added to the extracellular bathing solution, as indicated. (**B**) Amiloride-sensitive current before (“-”) and after (“+”) acid removal —circles show individual data points, lines (red) indicate means and SEMs (*n* = 15, \**P* < 0.0001 by paired 2-tailed Student’s *t* test). (**C**) Current-voltage relationship of amiloride-sensitive current before and after acid exposure (mean ± S.E.M, *n* = 4, most error bars are hidden by the data symbols). (**D**) Increase in amiloride-sensitive current (relative to pH 2 response) versus pH (mean ± S.E.M, *n* = 5). (**E**) Current-voltage families in HEK 293T cells expressing human αβ γENaC, obtained by whole cell patch clamp before (top) and after (bottom) acid exposure (pH 2.5), and in the absence (left) or presence (right) of amiloride (10 μM). (**F**) Amiloride-sensitive whole-cell conductance before and after acid exposure (*n* = 12, \**P* < 0.003 by paired 2-tailed Student’s *t* test).

Although acid stimulated ENaC, it did not disrupt characteristic biophysical properties of the channel—it’s linear current-voltage relationship (Fig. 1C), cation selectivity sequence (Li^+^>Na^+^>>K^+^ (Table 1)), and block by amiloride (Fig. 1A and Table 1) were intact. To determine the pH required to stimulate ENaC, we varied extracellular pH and quantified the increase in current after removing the acid (Fig. 1D). pH 3.0 did not increase ENaC current, but there was a small increase at pH 2.75, and maximal stimulation occurred at pH 2.25 (pH_50_ = 2.6).

**Table 1.** Effects of acid on ENaC biophysical properties.

|  | - Acid | + Acid | *n* | *P* value |
| --- | --- | --- | --- | --- |
| Li<sup>+</sup>/Na<sup>+</sup> permeability ratio | 1.41 ± 0.045 | 1.43 ± 0.018 | 4 | 0.74 |
| K<sup>+</sup>/Na<sup>+</sup> permeability ratio | 0.033 ± 0.007 | 0.042 ± 0.004 | 4 | 0.11 |
| Amiloride block (IC50) | 259 ± 42 nM | 202 ± 25 nM | 6 | 0.10 |
| Single channel conductance | 7.39 ± 0.22 pS | 6.13 ± 0.16 pS | 14-18 | 5 x 10<sup>-5</sup> |

We tested whether acid stimulation was dependent on cell type. In HEK 293 cells, we quantified ENaC whole-cell conductance by measuring current at voltages from -100 to +30 mV in the presence and absence of amiloride (Fig. 1, E and F). Exposure and washout of pH 2.5 increased ENaC conductance increased 2.6-fold (Fig. 1F), similar to results in *Xenopus* oocytes. However, the effect of acid was species-specific; in contrast to its stimulation of human ENaC, acid failed to stimulate mouse ENaC (Fig. S1).

Although acid stimulated ENaC, this stimulation was not evident until the acid was removed from the extracellular bathing medium (Fig. 1A). This finding raised the possibility that acid might have more than one effect on ENaC. To test this possibility, we stimulated ENaC with acid, removed the acid, then exposed it to acid (pH 2.5) a second time. Acid re-exposure decreased ENaC current, an effect that was rapidly and completely reversed upon acid removal (Fig. S2). Thus, acid in this pH range had two independent effects on ENaC; it irreversibly stimulated ENaC (stimulation was sustained in the absence of acid) and it reversibly inhibited ENaC. Because the two effects oppose one another, ENaC stimulation was only observed when the acid was removed from the extracellular solution. These opposing effects may be responsible in part for previous conflicting reports on whether acid modulates salt taste (*13-18*).

We investigated the mechanism by which acid stimulates ENaC by recording single channel currents before and after exposure to acid (pH 2.5). Fig. 2A shows representative current recordings—downward deflections indicate channel openings. Acid had two effects on ENaC current. First, it produced a small decrease in current amplitude, as demonstrated by the small shift in the single channel current-voltage relationship (Fig. 2B) and decrease in single channel conductance (Table 1). Second, acid dramatically increased the fraction of time that ENaC was in the open state (Fig. 2, A and C). This resulted from two effects on channel gating—an increase in mean open time (Fig. 2D) and a decrease in mean closed time (Fig. 2E). Thus, acid stimulates ENaC by enhancing channel opening and reducing channel closing.

**Fig. 2.**
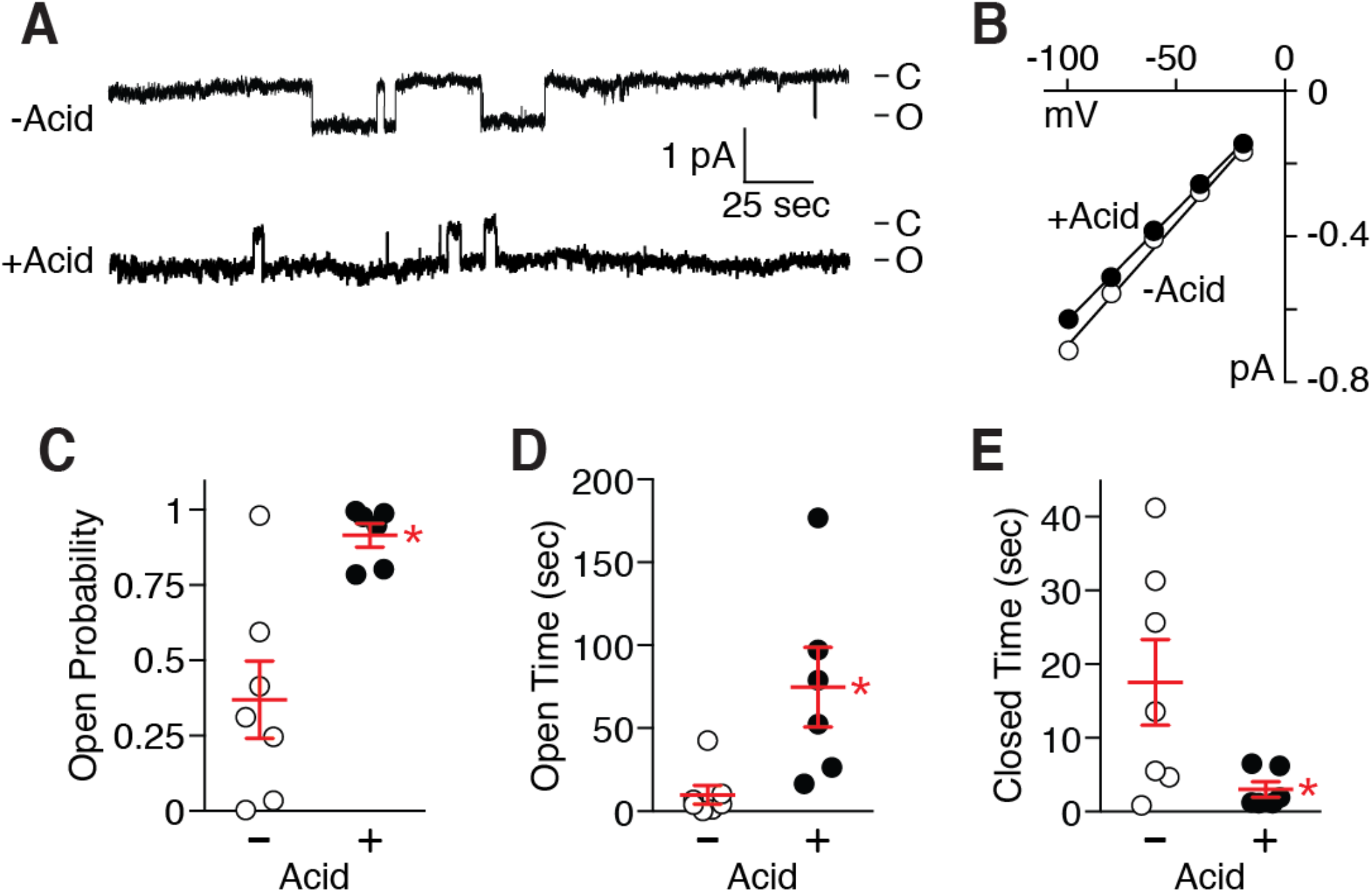
Acid increases ENaC open probability. (**A**) Single channel current traces (cell-attached configuration at -100 mV) from *Xenopus* oocytes expressing human αβ γENaC without (top) or with (bottom) acid pretreatment (pH 2.5). (**B**) Single channel current-voltage relationships from patches without or with acid pretreatment (mean ± SEM, *n* = 14-18, error bars hidden by data symbols). (**C**) Open probability determined from patches containing only one observed channel without (“-”) and with (“+”) acid pretreatment—circles show individual data points, lines (red) indicate means and SEM (*n* = 6-7, \**P* < 0.005 by unpaired 2-tailed Student’s *t* test). Mean open time (**D**) and closed time (**E**) were determined from these patches (\**P* < 0.02 and < 0.05, respectively, by unpaired 2-tailed Student’s *t* test).

Humans commonly consume highly acidic foods and beverages. We asked whether one such beverage, Coca-Cola® (pH 2.2), would stimulate ENaC. We replaced the extracellular bath solution with Coca-Cola® (supplemented with 110 mM NaCl to allow measurement of sodium current). This produced a small transient current increase followed by a decrease in current.

When we washed out the Coca-Cola® with pH 7.4 solution, ENaC current increased to a level 1.6-fold higher than current prior to Coca-Cola® exposure (Fig. 3, A and B). Pepsi® (pH 2.4) also stimulated ENaC (Fig. S3). Diet Coke® (pH 3.2) is less acidic than Coca-Cola®—it failed to increase ENaC current (Fig. 3B), consistent with our observation that stimulation requires pH more acidic than 3 (Fig. 1C).

**Fig. 3.**
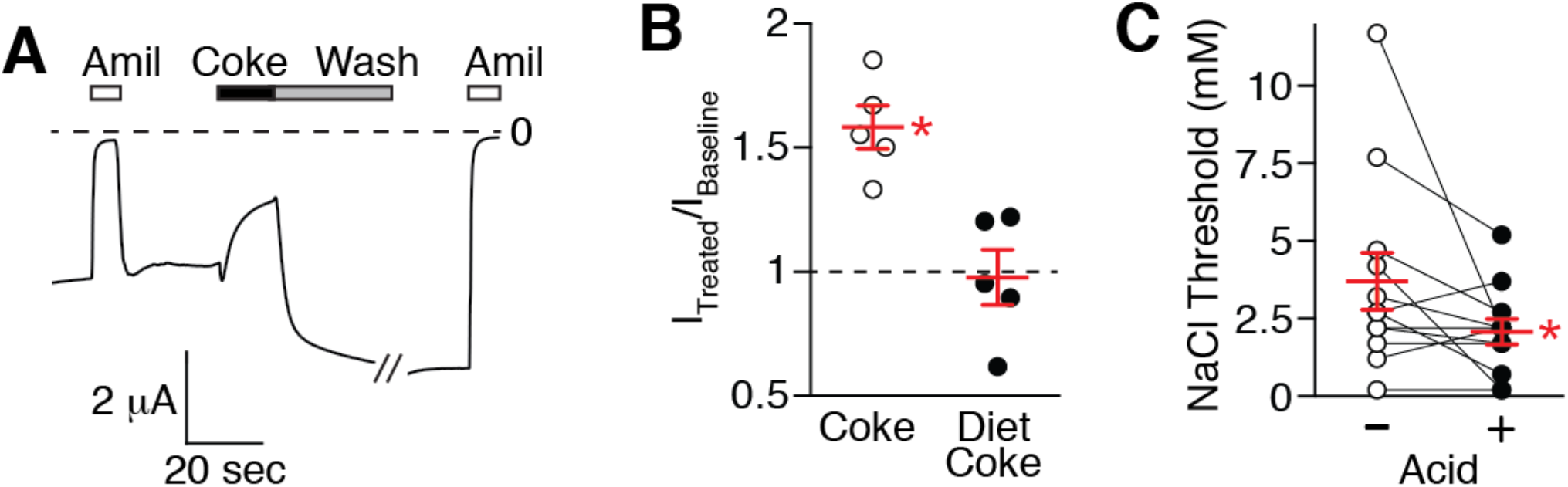
Acid increases salt taste. (**A**) Current trace from *Xenopus* oocyte expressing human αβ γENaC. Amiloride (10 μM) and Coca-Cola® (with 110 mM NaCl) were added to the extracellular bathing solution, as indicated. (**B**) Amiloride-sensitive current after treatment with Coca-Cola® or Diet Coke®, relative to baseline current before treatment—individual data points (circles) and mean/SEM (red lines) are shown (n = 5, \**P* < 0.002 versus baseline by paired 2-tailed Student’s *t* test). (**C**) Salt taste threshold in human subjects before (“-”) and after (“+”) exposing the tongue to 10 mM phosphoric acid (pH 2.2)—individual data points (circles) and mean/SEM (red lines) are shown (n = 12, \**P* < 0.04 by paired 1-tailed Student’s *t* test).

Based on these results and the observation that ENaC functions as a salt taste receptor (*5*), we predicted that acid would enhance salt taste in humans. To test this prediction, we asked normal volunteers to taste water samples containing varying concentrations of NaCl. The detection threshold for NaCl varied considerably between subjects (Fig. 3C), consistent with previous reports (*22-24*). However, exposing the anterior portion of the tongue to pH 2.2 for 15 sec reduced the NaCl detection threshold by almost half. Thus, following acid exposure, subjects were able to detect lower concentrations of NaCl.

We conclude that by increasing ENaC activity, acid enhances salt taste. Our findings highlight the complex interplay between different taste modalities; although salt and acid tastes are mediated by different receptors, these stimuli intersect at the level of ENaC. This provides a molecular explanation for the interaction between acid and salt taste, and it may explain the appeal of consuming salty foods, such as potato chips, in combination with acidic beverages, such as Coca-Cola® and Pepsi®. Moreover, our findings suggest that ENaC activation in the tongue could provide a strategy to alter sodium intake by enhancing the perceived saltiness of the foods we consume.

## Acknowledgments

We thank Diane Olson for technical assistance and Michael Welsh for advice and discussions. This work was supported by grants from the NIH (HL072256, to PMS; and 2T32HL007121-36, to DMC), the Department of Veterans Affairs Research Service (BX001862, to PMS; and BX000776, to CJB), and a pilot grant from The Institute for Clinical and Translational Science at the University of Iowa (U54TR001013), to PMS. The authors have no competing interests. All data are available in the manuscript or the supplementary materials.

## Supplementary Materials

Materials and Methods

Figures S1-S3

References 25

## Materials and Methods

### Electrophysiology

Currents were recorded in albino *Xenopus* laevis oocytes by two-electrode voltage clamp (holding potential -60 mV) one day after nuclear injection of cDNAs encoding human or mouse α-, β -, and γENaC (1 ng each), as described previously (*20*). The extracellular recording solution contained 116 mM NaCl, 2 mM KCl, 0.4 mM CaCl_2_, 1 mM MgCl_2_, 5 mM HEPES (pH adjusted to 7.4 with NaOH). For acidic solutions, pH was adjusted by addition of phosphoric acid. Coca-Cola® was shaken at room temperature to remove CO_2_ and 110 mM NaCl was added. ENaC current was quantified as the difference in current before and after addition of the ENaC blocker amiloride (10 μM) to the extracellular solution. For I/V relationships, current was stepped from - 60 mV to potentials from -100 and +80 mV (20 mV increments).

Single channel recordings (cell attached configuration) were obtained from wild type oocytes injected in the cytoplasm with cRNAs encoding human α-, β -, and γENaC (10 ng each). The pipet solution was 110 mM LiCl, 2 mM KCl, 1.54 mM CaCl_2_, 10 mM HEPES (pH 7.4 with LiOH), and bath solution was 110 mM LiCl, 2 mM KCl, 1.54 mM CaCl_2_, 10 mM HEPES, pH 7.4 with LiOH. Oocytes were exposed to acid (pH 2.5) or control (pH 7.4) bathing solution for 30 sec prior to recording. Single channel properties were quantified in patches containing a single channel using TAC (version 3.0.8; Bruxton Corporation), as described previously (*21*).

Whole cell patch clamp recordings were obtained from HEK 293T cells transfected with human α-, β -, and γENaC (2 μg each using Lipofectamine 2000, Invitrogen). The internal pipet solution was 100 mM NaCl, 10 mM EGTA, 5 mM MgCl_2_, 2 mM ATP, 0.3 mM GTP, 40 mM HEPES (pH 7.4 with KOH), and the external bathing solution was 120 mM NaCl, 5 mM KCl, 2 mM CaCl_2_, 1 mM MgCl_2_, 10 mM HEPES, 10 mM 2-(*N*-morpholino)ethanesulfonic acid (pH 7.4 with NaOH, and 2.5 with phosphoric acid). Voltage was stepped from -70 mV to voltages from -100 to +30 mV (10 mV increments).

### Sodium detection threshold

We determined sodium detection threshold using a three-choice, forced-choice, staircase assay (*25*) before and after exposing of the tip of the tongue to 10 mM phosphoric acid (pH 2.2) for 15 sec, followed by a rinse with deionized water. Subjects tasted three solutions presented in random order to identify which was different—two were deionized water, one was deionized water containing NaCl (0.2-15 mM). We began with 15 mM NaCl and reduced the NaCl concentration in subsequent trials. When an incorrect choice was made, we increased to a higher NaCl concentration. When the subject was unable to identify the NaCl solution three times, the NaCl threshold was calculated as the mean value of the lowest correctly identified NaCl solutions.

### Statistics

Student’s *t* test (1- or 2-tailed, paired or unpaired, as indicated in the figure legends) was used, with *P* < 0.05 considered significant.

### Study approval

The salt threshold protocol was approved by the Human Subjects Institutional Review Board at the University of Iowa. Subjects provided written informed consent prior to inclusion in the study. Animal protocols (*Xenopus laevis*) were approved by the University of Iowa IACUC and animal care was in accordance with institutional guidelines.

**Fig. S1.**
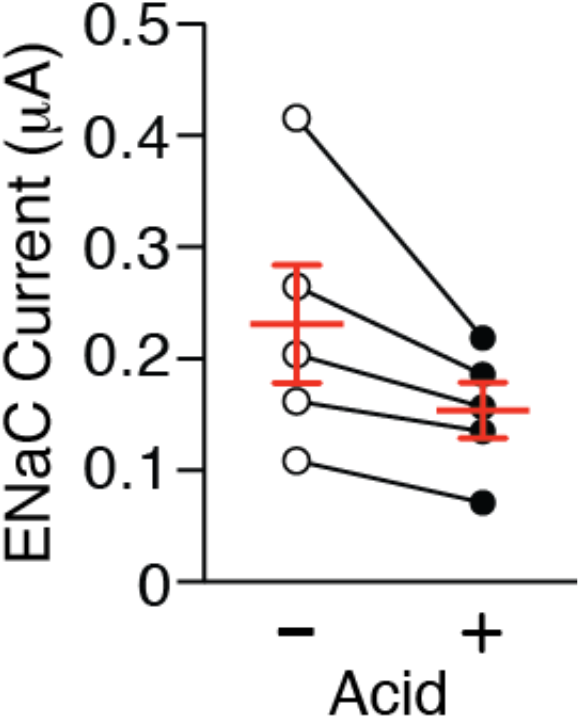
ENaC activation by acid is species specific. Amiloride-sensitive current before (“-”) and after (“+”) acid removal in *Xenopus* oocyte expressing mouse αβ γENaC obtained by two-electrode voltage clamp (holding potential -60 mV). Circles show individual data points, lines (red) indicate means and SEMs (*n* = 5, *P* = 0.067 by paired 2-tailed Student’s *t* test).

**Fig. S2.**
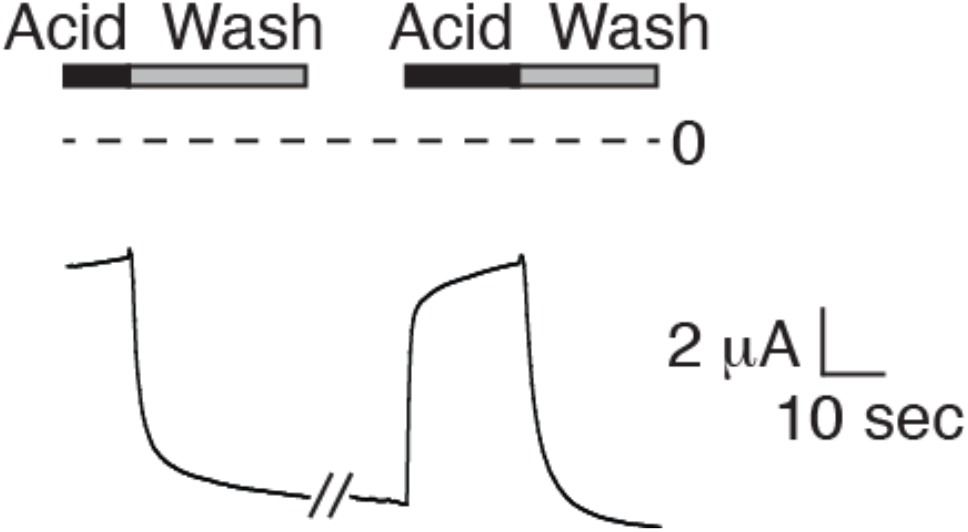
Acid reversibly inhibits ENaC. Current trace from a *Xenopus* oocyte expressing human αβ γENaC (holding potential -60 mV). Acid (pH 2.5) was added sequentially to the extracellular bathing solution and washed out, as indicated.

**Fig. S3.**
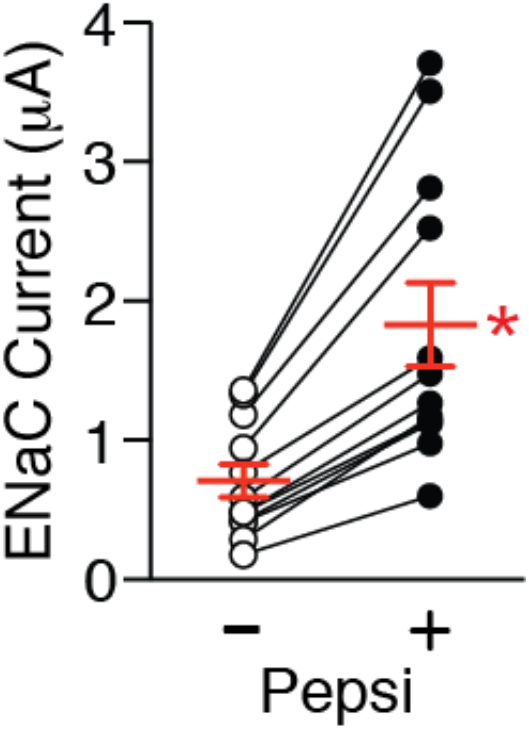
Pepsi stimulates ENaC current. Amiloride-sensitive current before (“-”) and after (“+”) acid removal in *Xenopus* oocyte expressing human αβ γENaC obtained by two-electrode voltage clamp (holding potential -60 mV). Circles show individual data points, lines (red) indicate means and SEMs (*n* = 12, \**P* < 0.0001 by paired 2-tailed Student’s *t* test).

